# Computational Disruption of Paired Helical Filaments (PHFs) Assembly Using Milk Lactalbumin-derived Peptides Against Alzheimer’s Disease

**DOI:** 10.1101/2024.12.30.630760

**Authors:** Saeed Pourmand, Sara Zareei, Mohanna Jozi, Shokoufeh Massahi

## Abstract

Peptides show great potential in diagnosing and treating Alzheimer’s disease (AD), particularly by targeting amyloid-beta plaques and neurofibrillary tangles (NFTs) formed by hyperphosphorylated tau proteins. This study focuses on designing peptide inhibitors from bovine milk alpha-lactalbumin to reduce tau aggregation in AD. Using computational techniques, such as docking, molecular dynamics simulations, and mutagenesis, we evaluated the binding and stability of these peptides against the paired helical filament (PHF) core. Our results identified promising inhibitors, with p136 emerging as the most effective. It significantly altered the PHF core’s structure, preventing further aggregation by blocking additional subunits. Additionally, p76 displayed strong binding against straight filaments (SFs). These findings highlight the potential of peptides derived from bovine milk alpha-lactalbumin as diagnostic and therapeutic tools for AD, with p136 standing out as a promising candidate for disrupting tau aggregation.

## 1. Introduction

Peptides have emerged as a promising tool in the diagnosis and treatment of many diseases such as cancer, viral infection, and Alzheimer’s disease (AD) [1-3]. In diagnostic applications, peptides-based probes and biomarkers have shown significant potential for early detection of AD. Among these, amyloid-beta (Aβ)-derived peptides are the most well-known peptide biomarkers, which have proven effective in diagnosing the onset of AD and in assessing the progression of the disease [4]. Peptides can also act as vectors or probes that can specifically bind to AD biomarkers and assist in the early detection of AD especially through imaging techniques like positron emission tomography (PET) or magnetic resonance imaging (MRI) [5]. For Example, D-enantiomeric peptides have been developed as molecular probes to detect Aβ1-42 plaques in the living brain at submicromolar concentrations [6, 7].

Peptides are been explored for their therapeutic potential in AD, particularly as inhibitors targeting various pathological pathways involved in the disease. A major focus has been on preventing amyloid-beta (Aβ) aggregation, with peptide inhibitors designed to block the formation, elongation, or aggregation of amyloid plaques and their toxic oligomers [8, 9]. Beyond amyloid, peptide inhibitors have been developed to target other critical molecular contributors to AD pathogenesis, such as β-Site APP Cleaving Enzyme 1 (BACE1), which drives amyloid production; Glyceraldehyde-3-Phosphate Dehydrogenase (GAPDH), associated with apoptotic cell death, [10]; Tyrosine Phosphatase (TP), a regulator of tau phosphorylation [11]; and the potassium channel KV1.3, which affects neuroinflammation [12].

Neurofibrillary tangles (NFTs) are composed of abnormal intracellular aggregates of hyperphosphorylated tau proteins, which are another hallmark of neurodegenerative disorders, including AD [13, 14]. Under normal conditions, tau proteins stabilize microtubules, particularly in the distal parts of neurons, ensuring proper transport of materials within neurons. However, in pathological states, hyperphosphorylated tau detaches from microtubules and aggregates into NFTs [15], disrupting cellular functions and contributing to neurodegeneration. These tau deposits manifest in two distinct morphological structures: paired helical filaments (PHFs) and straight filaments (SFs), with frequencies of 95% and 5% in AD, respectively [16]. Both structures consist of double helical stacks of C-shaped subunits, although they differ in topology. The core region of PHF (297-391) plays a crucial role in tau aggregation and is highly neurotoxic, making it a promising target for anti-Alzheimer’s therapies [17].

Recent research has also explored the application of milk and its components in the diagnosis and treatment of Alzheimer’s disease, driven by findings that consistent diary intake may reduce the risk of dementia [18-21]. Studies have shown that goat milk possesses neuroprotective properties and can ameliorate memory deficits in both mouse models and human patients. These benefits are attributed to its ability to increase glutathione levels and decrease malondialdehyde, acetylcholinesterase, and serum total cholesterol activity [22]. Additionally, certain milk-derived peptides, such as βb-casomorphin-5 and βb-casomorphin-7, have shown potential in treating AD by inhibiting acetylcholine aceterase activity, a key enzyme involved in the degradation of acetylcholine, which is critical for cognitive function [23].

In this study, we used bovine milk alpha-lactalbumin as a template for rationally designing peptides, and we assessed their anti-Alzheimer activity through the inhibition of PHF aggregation. By focusing on these novel peptides, we aim to contribute to the development of more effective diagnostic and therapeutic strategies for AD.

## 2. Computational Approaches

### 1. Template and Target Preparation and Initial Docking

The full length of apo-bovine alpha-lactalbumin (α-Lac) was extracted from the RCSB databank under the accession number 1F6R [24]. Next, chain A of this hexamer was chosen and docked against the A, C, E, G, and I of PHF under the accession number of 5O3L [25] using the HADDOCK server (https://rascar.science.uu.nl/) and giving the interface between C-shaped subunits (residues 331-338). The lactalbumin was allowed to accommodate freely to this binding region. The docking of straight filaments (SFs) was also carried out using the PDB ID: 5O3T [25]. Each peptide was generated by shifting one residue at a time along the full sequence, resulting in a comprehensive library of 15-mer peptides.

### 2. Molecular Dynamics (MD) Simulation

The simulations were conducted using GROMACS package version 2019. Topology and coordinates files were generated using CHARMM27 forcefield. Each complex was solvated in a simple point-charge (SPC) water box with periodic boundary conditions, and chloride ions were added to neutralize the systems. A total of 22 Cl^−^ ions were added to the P67 and P76 systems, while 23 Cl^−^ ions were required for the template and P136 systems. The apo-protein system required 25 Cl^−^ ions to achieve neutrality.

Next, the system underwent energy minimization of 50,000 steps to remove steric clashes and relax the structure. Then, the minimized system was then equilibrated under NVT (constant number of particles, volume, and temperature) at a constant temperature of 300K and NPT (constant number of particles, pressure, and temperature) conditions until the reaching 1 bar pressure. The production phase of the 150 ns molecular dynamics (MD) simulation employed the Particle Mesh Ewald (PME) method to account for long-range electrostatic interactions, with the Verlet algorithm used to calculate the trajectories.

To analyze the resulting data, various metrics were computed, including root-mean-square deviation (RMSD), root-mean-square fluctuation (RMSF), radius of gyration (Rg), solvent-accessible surface area (SASA), hydrogen bond analysis, and Molecular Mechanics Poisson-Boltzmann Surface Area (MM-PBSA) calculations. Principal Component Analysis (PCA) was also conducted using gmx covar and gmx anaeig, enabling the generation of a covariance matrix, diagonalization, and projections onto principal components. The trajectory was saved at regular intervals for further analysis.

### 3. Mutagenesis

We used the MCSM (Membrane Curvature Server for Molecular Dynamics) server was employed to introduce mutations in peptide sequence based on the Delta Delta G (ΔΔG) concept. This approach allows for the prediction of stability changes in peptides caused by single-point mutations. The ΔΔG value represents the difference in binding free energy between the mutant peptide and the wild-type peptide, calculated using the formula:

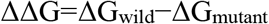

In this context, positive ΔΔG values indicate that the mutation confers a stabilizing effect on the peptide, enhancing its overall stability. Conversely, negative ΔΔG values suggest that the mutation destabilizes the peptide, reducing stability. Additionally, ΔΔG values close to zero indicate mutations that are virtually neutral concerning their impact on peptide stability, showing no significant effect.

To facilitate the generation of diverse mutated peptide sequences, we employed the MutLib.py script (https://github.com/SaraZareei/Peptide-Design/blob/main/MutLib.py). This script accepts all favorable residues for mutation and generates all possible sequence combinations.

## 3. Results and Discussion

### Structural Analysis of PHF Core and Template Selection

The Cryo-EM structure of the PHFs core, extracted from a post-mortem Alzheimer’s cerebral cortex included two protofilaments of tau protein between residues 306-378. The interaction forming the interface between these protofilaments was between residues 331-338 of each strand. This region was considered a target against the assembly of tau aggregates where the lactalbumin was docked against it. The docking results revealed that the lactalbumin residues Glu1, Gln2, Leu3, Thr4, Lys5, Phe31, His32, Thr33, Ser34, Gly35, Asn37, Gln39, Ala40, Ile41, Val42, Gln43, Asn44, Asn45, Glu49, Gln54, Leu110, Lys114, and Gln117 interacted with PHF residues with HADDOCK score of 17. Out of them, we extracted residues 21-45 as a template and docked it against the interface and re-docked against the target. The results showed that the template peptide bound to the PHF core by a combination of hydrogen bonds and hydrophobic interactions with a binding score of −79.8 (Figure 1).

**Figure 1.**
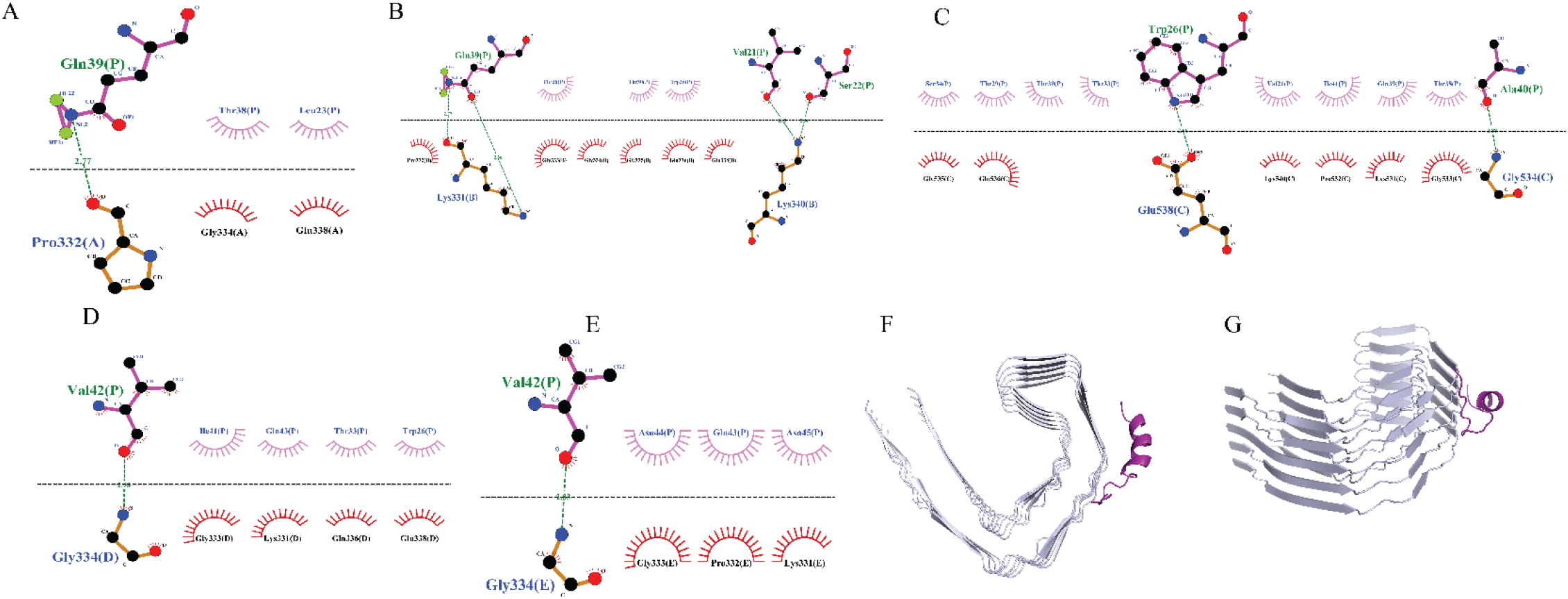
The docking results of template peptide with PHF core in 2D (A-E) and 3D (F and G) presentations. The template is shown in violet while the PHF core is depicted in silver in the last two pictures.

### Peptide Library

To enhance the binding potential of the peptide inhibitor, we used MMPBSA analysis following a 150ns MD simulation. The results showed the contribution of each template peptide residue to the overall interaction with the PHF core (Table 1). MMPBSA calculation identified Val21, Thr29, His32, Ser34, Gly35, Ala40, and Val42 with positive binding free energy contributions, indicating unfavorable interactions. These residues were considered hotspots for mutagenesis. MCSM output evaluated the impact of single mutations of all twenty standard amino acids on the stability of the template (Supplementary Table S1). The positive ΔΔG values indicate the stabilizing point mutations (Table 2).

**Table 1.** Energy Decomposition of Template Residues Using MMPBSA Analysis.

|  |  |
| --- | --- |
| Residues | Total Energy |
| Val-21 | 147.01 |
| Ser-22 | -5.08 |
| Leu-23 | -13.26 |
| Pro-24 | -3.80 |
| Glu-25 | -139.59 |
| Trp-26 | -20.44 |
| Val-27 | -2.19 |
| Cys-28 | -2.54 |
| Thr-29 | 0.58 |
| Thr-30 | -4.22 |
| Phe-31 | -1.45 |
| His-32 | 0.89 |
| Thr-33 | -2.79 |
| Ser-34 | 0.66 |
| Gly-35 | 4.53 |
| Tyr-36 | -7.93 |
| Asp-37 | -207.70 |
| Thr-38 | -1.10 |
| Gln-39 | -2.99 |
| Ala-40 | 0.98 |
| Ile-41 | -3.98 |
| Val-42 | 1.82 |
| Gln-43 | -1.47 |
| Asn-44 | -1.86 |
| Asn-45 | -202.60 |

**Table 2.** Stabilizing Point Mutations on Template Sequence Using MCSM Server.

|  |  |  |  |
| --- | --- | --- | --- |
| Position | Wild Residue | Mutated Residue | Predicted $\Delta\Delta G$ |
| 29 | T | M | 0.378 |
| 32 | H | A | 0.069 |
| 32 | H | V | 0.504 |
| 32 | H | L | 0.684 |
| 32 | H | S | 0.016 |
| 32 | H | W | 0.495 |
| 32 | H | T | 0.133 |
| 32 | H | Q | 0.196 |
| 32 | H | E | 0.257 |
| 32 | H | C | 0.3 |
| 32 | H | P | 0.504 |
| 32 | H | D | 0.241 |
| 32 | H | F | 0.492 |
| 32 | H | I | 0.684 |
| 32 | H | N | 0.139 |
| 32 | H | M | 0.536 |
| 32 | H | Y | 0.836 |
| 32 | H | K | 0.095 |
| 34 | S | V | 0.021 |
| 34 | S | L | 0.03 |
| 34 | S | P | 0.021 |
| 34 | S | I | 0.03 |
| 34 | S | M | 0.459 |
| 35 | G | D | 0.006 |
| 42 | V | R | 0.29 |
| 42 | V | N | 0.035 |

Using these data, we generated a library of 170 peptides out of which 38 peptides were identified as non-allergen and non-toxic and next docked against the PHF core (Supplementary table S2, Table 3). The docking results showed that p67, p76, and p136 had the lowest binding scores of −111.5, −109.2, and −113.2, respectively, indicating the highest potential for binding to the PHF core.

**Table 3.**
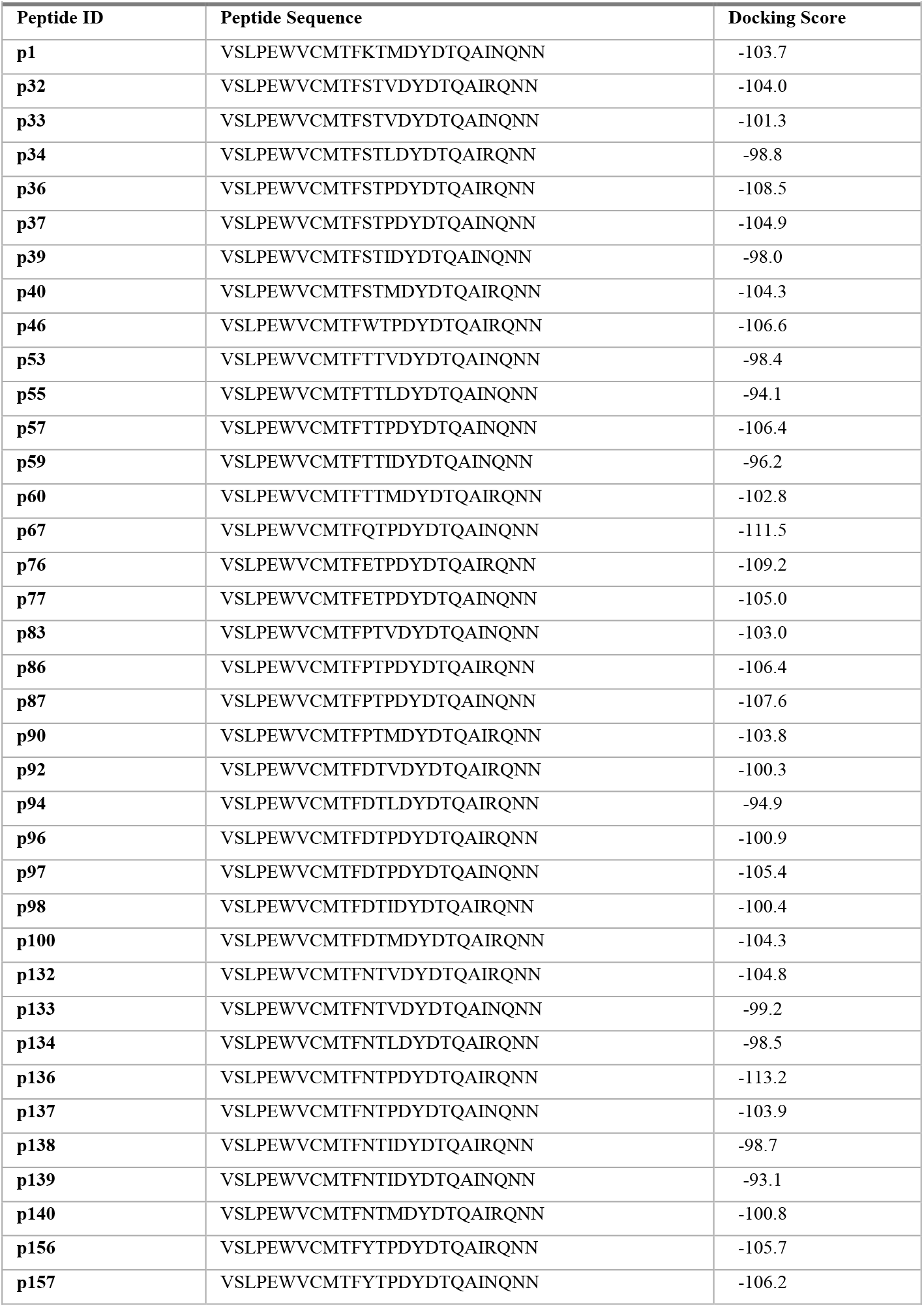
The final library of mutated non-allergen non-toxic peptides and their docking scores.

The analysis of the interactions showed that the designed peptides attached to the PHF core with both hydrogen bonds and hydrophobic interactions (Table 4, and Figures S1, S2, and S3). P67 showed the lowest number of H-bonds and hydrophobic interaction with the target, explaining its lowest binding score. Although p76 showed the same number of hydrogen bonds as p136 (twenty interactions), it exhibited a slightly lower number of h-bond involvement, indicating the underlying reason for the higher binding affinity of p136. It can be concluded that chains B and C are the most influential in the potential inhibitory activity of the peptides due to the highest number of interactions they are involved in. Chains D and A also play important roles, while Chain E, is less critical compared to the others.

**Table 4.**
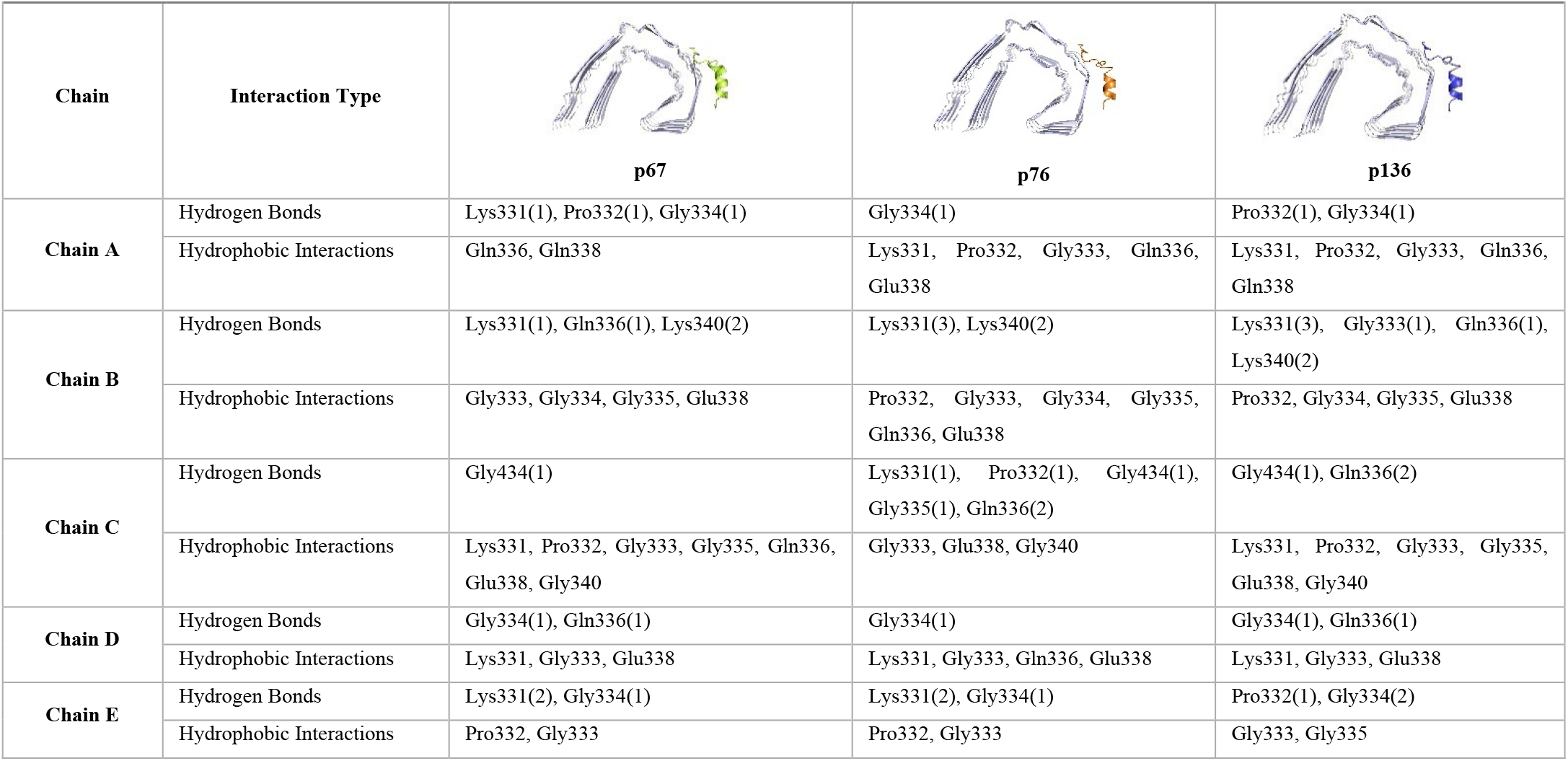
The profile of interactions between PHF core and peptides p67, p76, and p136. Parentheses indicate the number of hydrogen bonds while the other residues indicate hydrophobic interactions.

| Chain | Interaction Type | p67 | p76 | p136 |
| --- | --- | --- | --- | --- |
| Chain A | Hydrogen Bonds | Lys331(1), Pro332(1), Gly334(1) | Gly334(1) | Pro332(1), Gly334(1) |
|  | Hydrophobic Interactions | Gln336, Gln338 | Lys331, Pro332, Gly333, Gln336, Glu338 | Lys331, Pro332, Gly333, Gln336, Gln338 |
| Chain B | Hydrogen Bonds | Lys331(1), Gln336(1), Lys340(2) | Lys331(3), Lys340(2) | Lys331(3), Gly333(1), Gln336(1), Lys340(2) |
|  | Hydrophobic Interactions | Gly333, Gly334, Gly335, Glu338 | Pro332, Gly333, Gly334, Gly335, Gln336, Glu338 | Pro332, Gly334, Gly335, Glu338 |
| Chain C | Hydrogen Bonds | Gly434(1) | Lys331(1), Pro332(1), Gly434(1), Gly335(1), Gln336(2) | Gly434(1), Gln336(2) |
|  | Hydrophobic Interactions | Lys331, Pro332, Gly333, Gly335, Gln336, Glu338, Gly340 | Gly333, Glu338, Gly340 | Lys331, Pro332, Gly333, Gly335, Glu338, Gly340 |
| Chain D | Hydrogen Bonds | Gly334(1), Gln336(1) | Gly334(1) | Gly334(1), Gln336(1) |
|  | Hydrophobic Interactions | Lys331, Gly333, Glu338 | Lys331, Gly333, Gln336, Glu338 | Lys331, Gly333, Glu338 |
| Chain E | Hydrogen Bonds | Lys331(2), Gly334(1) | Lys331(2), Gly334(1) | Pro332(1), Gly334(2) |
|  | Hydrophobic Interactions | Pro332, Gly333 | Pro332, Gly333 | Gly333, Gly335 |

Following the docking procedure, the most potent peptide inhibitors were subjected to 300 ns of MD simulations to investigate their impact on the conformation PHF core. The RMSD results indicated that the peptides p67 and p76 (with the average RMSD values of 0.54 nm and 0.49 nm, respectively) reduced RMSD in the PHF core (average RMSD = 0.85nm), suggesting that these peptides bind in a manner that preserves the overall structure of the PHF core (Figure 1-A). This minimal deviation implies a stable interaction that does not significantly alter the core’s native conformation.

In contrast, the peptide p136 induced substantial conformational changes in the PHF core (RMSD _average_=67 nm), showing a marked divergence from the apo form (unbound PHF core) (Figure 2A). This significant deviation could suggest a more disruptive binding mechanism, which may interfere with the normal assembly process of the PHF. Such disruption could be advantageous for inhibiting the formation of PHF assemblies, highlighting p136 as a promising inhibitor with the potential to alter the structural dynamics of the PHF core more effectively than p67 and p76.

**Figure 2.**
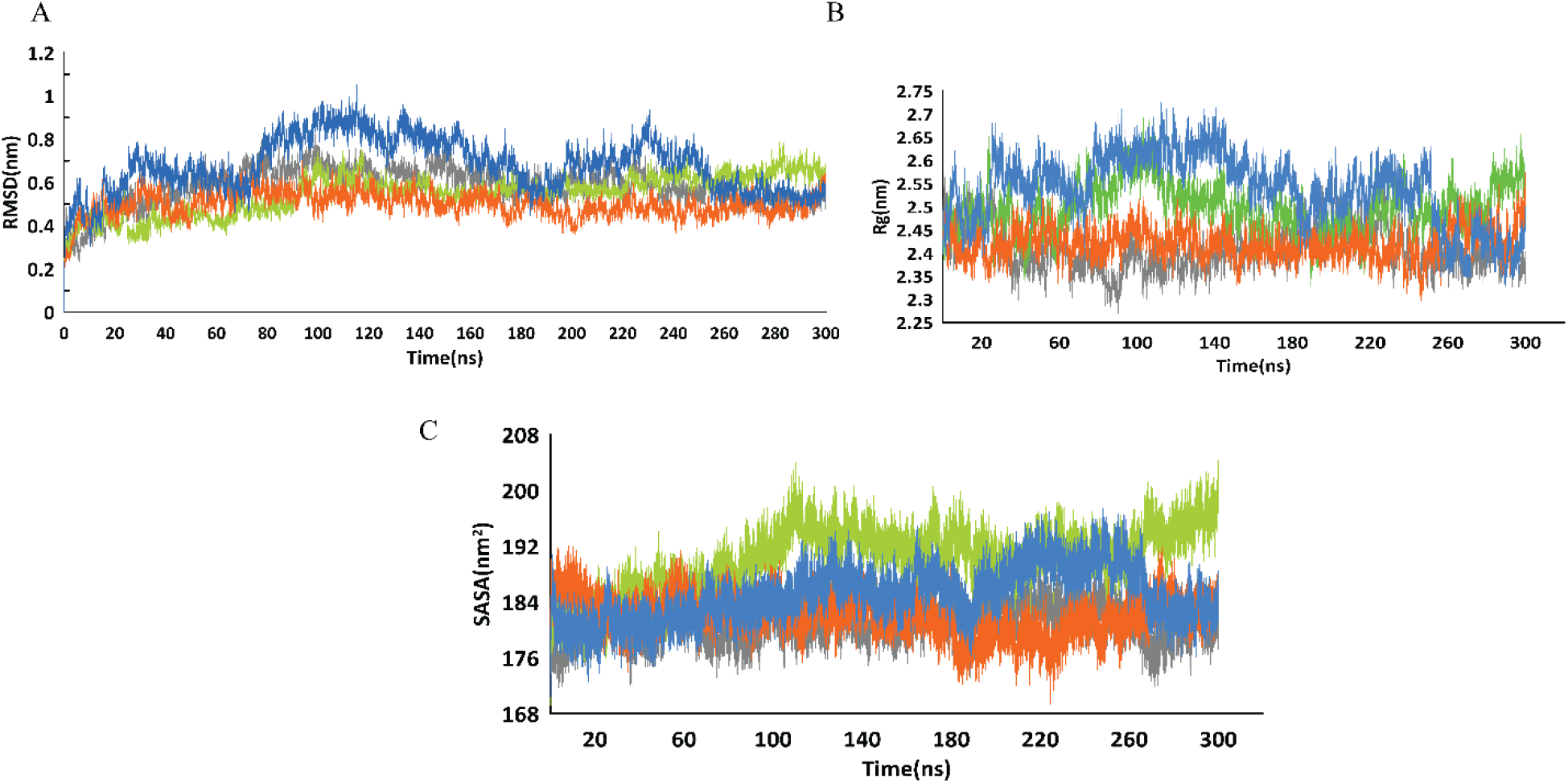
MD results of PHF core (grey) in complex with p67 (limon), p76 (orange), and p136 (blue). The plots show RMSD (A), Rg (B), and SASA (C) results for each complex.

Further analysis using the Rg revealed that all three peptide inhibitors increased the Rg of the PHF core (with the average Rg of 2.49, 2.42, and 2.53 for p67, p76, and p136, respectively compared with average value of 2.40 for PHF core), indicating that the protein adopted a more extended conformation (Figure 2B). This suggests that the PHF subunits transitioned from their native assembly to a more dispersed spatial distribution. Among the peptides, p136 caused the most significant increase in Rg, indicating that its binding induced a greater separation between subunits compared to the other peptides.

In a similar manner, the average SASA results were 190.2, 182.0 and 184.9 nm for p67, p76, and p136, respectively compared to 181.2 for PHF core. It can be inferred that the peptides may induce domain movements that rearrange the domains relative to each other, leading to a more dispersed structure (Figure 2C).

As inhibitors of PHF assembly, peptides should ideally increase the flexibility and mobility of the residues involved in the assembly. Since nearly all residues in each chain participate in the stacked conformation of PHF chains, monitoring the mobility of all residues is crucial. RMSF analysis revealed that peptide binding resulted in increased flexibility and destabilization of residues across each chain. This effect was particularly pronounced with peptides p136 and p67 (Figure 3).

**Figure 3.**
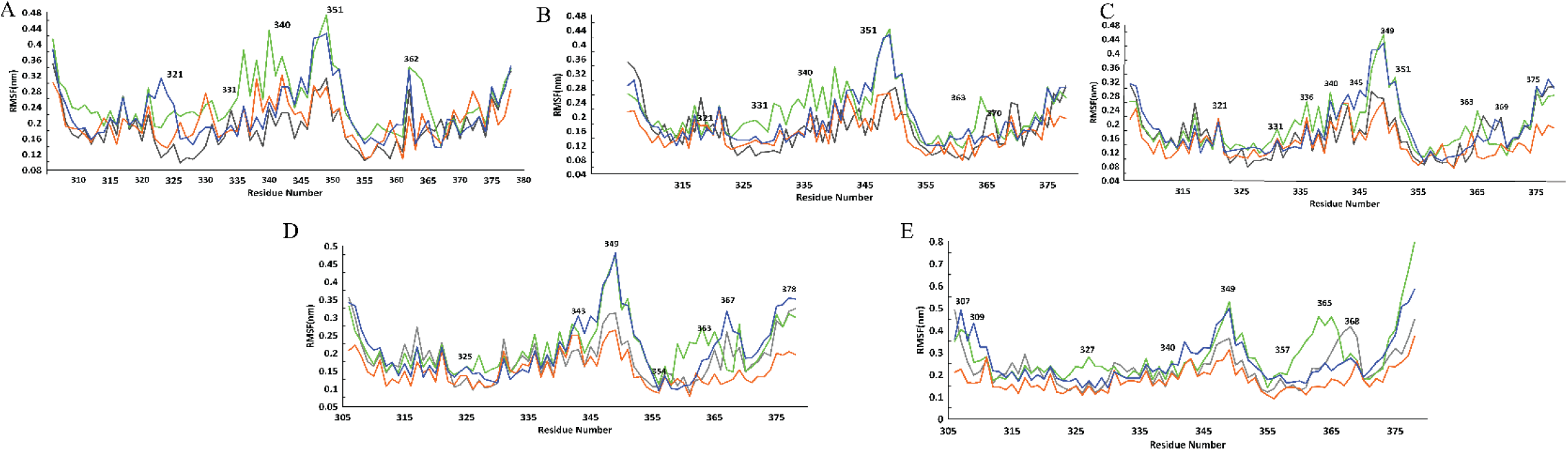
RMSF plots of peptides p67 (lime), p76 (orange), and p136 (blue), compared with the PHF core in its apo form (gray) for monomers A, B, C, D, and E.

Another critical factor supporting the peptide inhibitors’ ability to disrupt the assembly and aggregation of the PHF core is the reduction in the number of contacts between the PHF core chains. Molecular dynamics analysis revealed that the total number of interchain molecular contacts within the PHF core significantly decreased in the presence of the designed peptides. This finding indicates that the peptides interfered with the molecular organization and assembly of the PHF core, reducing its propensity to form the ordered filaments characteristic of Alzheimer’s disease (Figure 4). However, the number of contacts between the peptides and the PHF core remained relatively stable, with no significant decrease observed in the contact count (Supplementary Figure S4).

**Figure 4.**
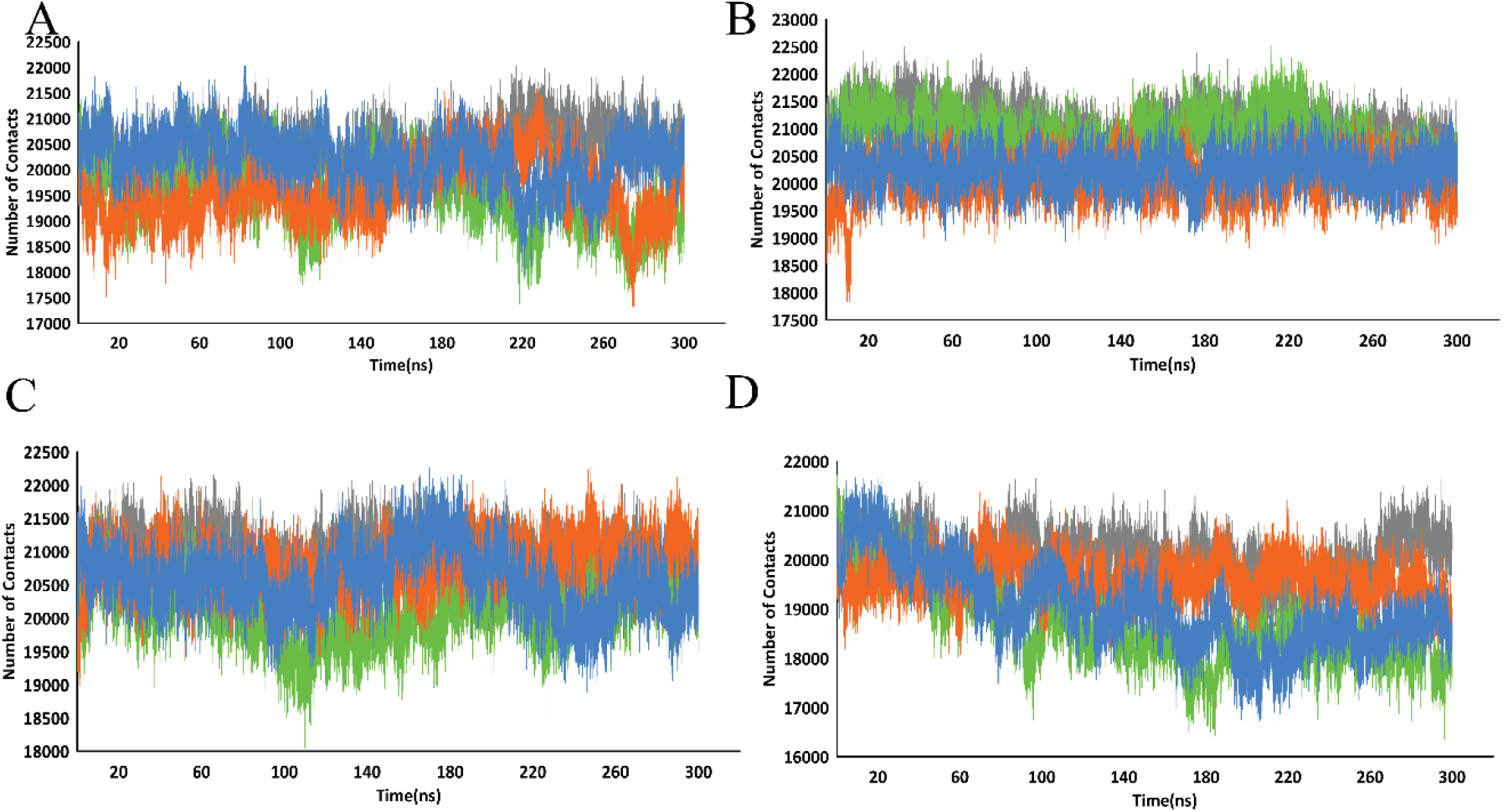
Total number of contacts between PHF chains A-B (A), B-C (B), C-D (C), and D-E (D) in the presence of the designed peptides p67 (green), p76 (orange), and p136 (blue) compared to the apo PHF core (gray).

Additionally, the MD simulations revealed that the designed peptides introduced variability in the distances between the chains, resulting in an irregular structure. The peptides increased the separation between chain pairs compared to the tightly assembled structure of the PHF core. This suggests that the peptide inhibitors prevent the chains from coming into proximity and interacting with each other, potentially leading to disassembly or structural distortions within the filaments (Figure 5). This is comparable to the stable distance between the peptides and the PHF core, suggesting that the peptides may not detach from the PHF core until a sufficiently disruptive distance between the chains is reached (Supplementary Figure S4-B).

**Figure 5.**
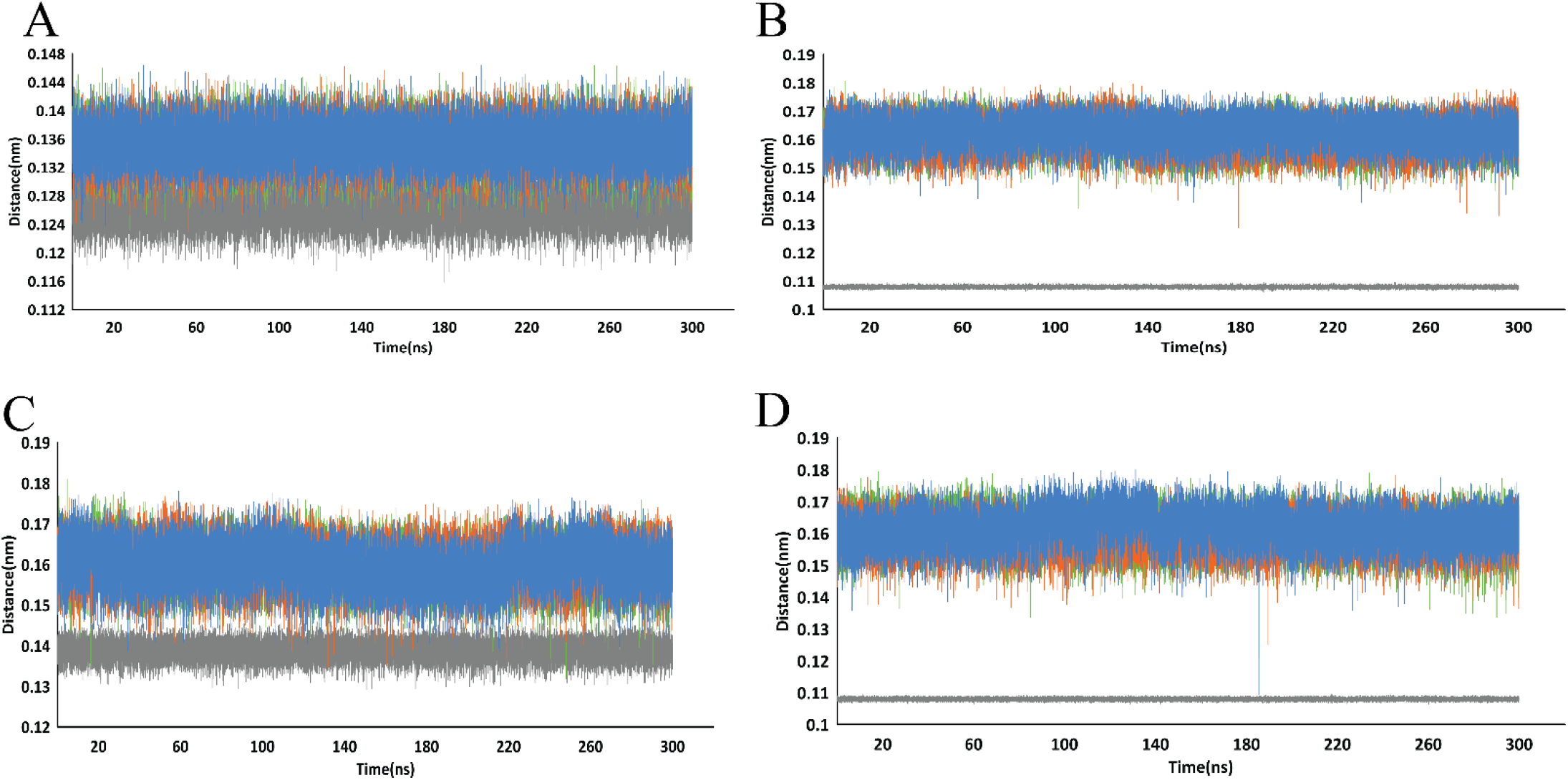
Variability in distances between PHF chains A-B (A), B-C (B), C-D (C), and D-E (D) in the presence of the designed peptides p67 (green), p76 (orange), and p136 (blue), compared to the apo PHF core (gray). The increased distances indicate that the peptides disrupt the close interactions between chains, leading to a more irregular and potentially destabilized PHF structure.

The impact of p67, p76, and p136 on the assembly of the PHF core is further highlighted by PCA analysis, which visually compares the conformational space explored by the PHF core before and after the addition of these peptide inhibitors (Figure 6). In the 2D projection of the last 10 ns of the PCA for the apo (unbound) PHF core, a wide conformational spread seen without any particular conformational cluster. This suggests that PHF core adopt a range of conformations. The projections of p67 and p76showed that these peptides induced two distinct conformational clusters in PHF having spaces sampled closely with PHF, showing that fewer conformations are stable upon the binding of P67 and p76.

**Figure 6.**
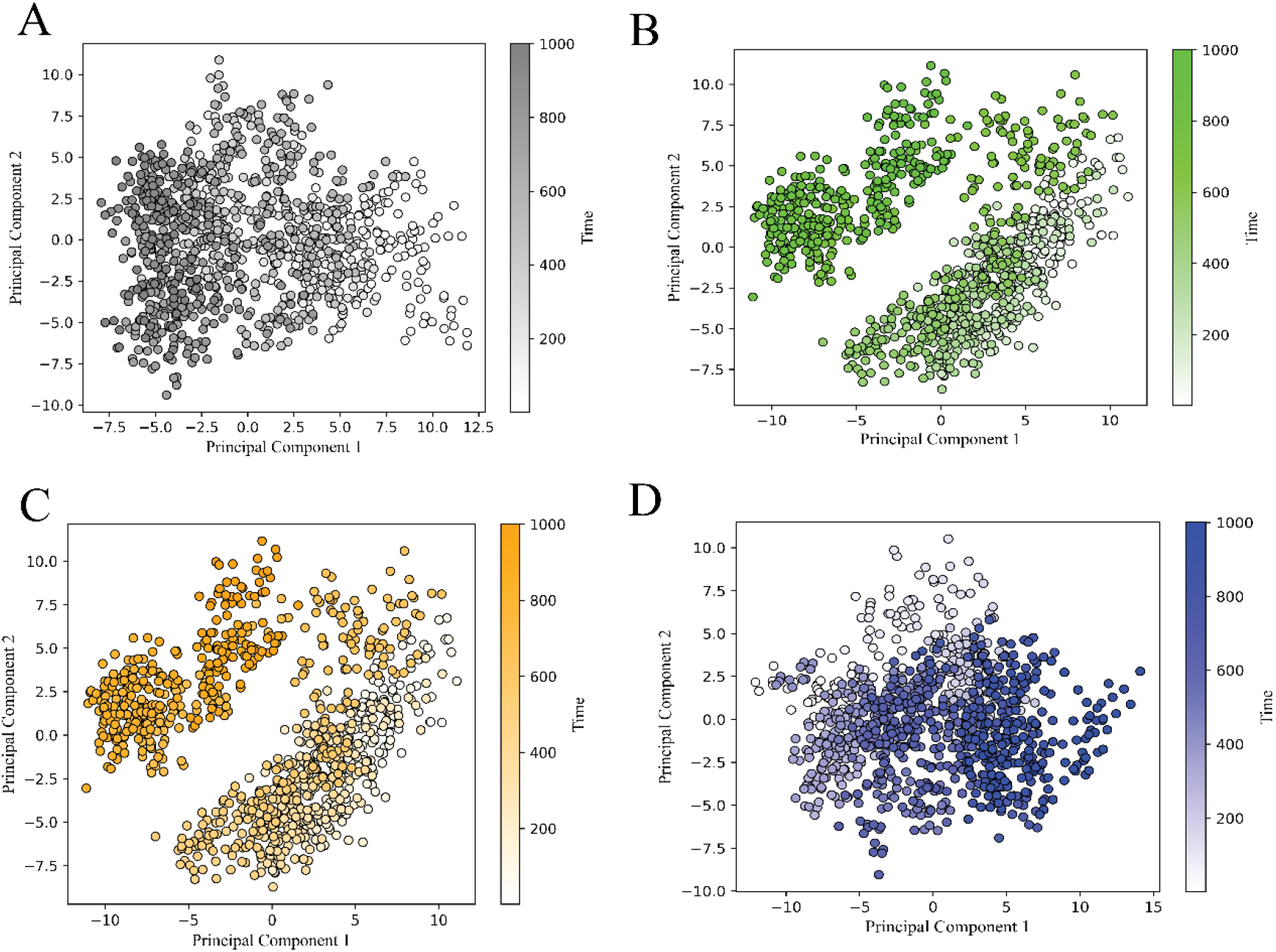
Principal Component Analysis (PCA) of the conformational changes in the PHF core over a 150 ns simulation period.

While the PCA plot for P136 may seem similar to the apo system, closer examination reveals that P136 induced a larger range of conformational changes along the PC1 axis (extending to 15, compared to 10 for the apo protein). This implies that P136 introduced significant conformational shifts along a dominant motion, potentially indicative of a larger structural rearrangement.

Comparing the 2D projections of the peptides, it can be concluded that p136 is the most effective in disrupting the PHF core assembly. Its greater dispersion and more distinct shift from the apo state indicate that p136 is particularly potent at destabilizing the PHF core.

To investigate whether these peptides can inhibit further aggregation by blocking the binding of additional PHF subunits, we performed docking studies of each PHF subunit against the already-formed peptide-PHF complex. The results demonstrated that p67, p76, and p136 effectively prevent the binding of the other PHF subunit, disturbing its ability to complete the aggregate complex. This inhibition likely contributes to the prevention of subunit aggregation (Figure 7).

**Figure 7.**
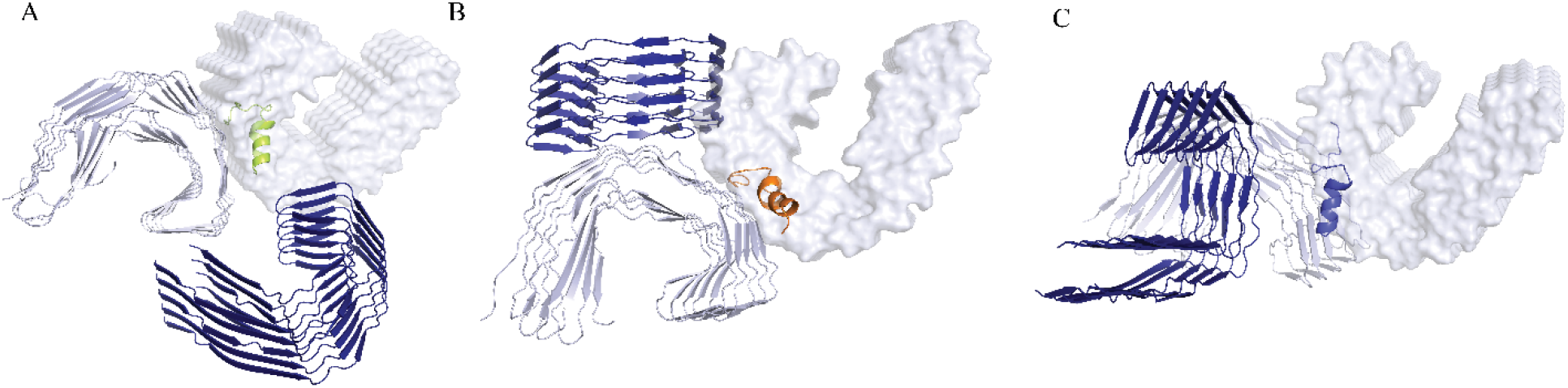
Control docking analysis of peptide-PHF complexes. The interaction between p67 (A, Limon), p76(B, orange), and p136(C, blue)-PHF complexes with additional PHF subunit (heavy blue). The expected coordinates of additional PHF subunits in PHF aggregates are shown in transparent silver.

We also checked the potential inhibitory activity of p67, p76, and p136 against SFs aggregation (Figure 7), which are formed by hyperphosphorylated tau [16] and are found in NFTs. Each peptide was docked against an SF subunit and demonstrated the ability to bind to SF monomers with binding scores of −80, −86, and −80, respectively. Notable, p76 showed the most potent binding ability against SFs (Figure 8).

**Figure 8.**
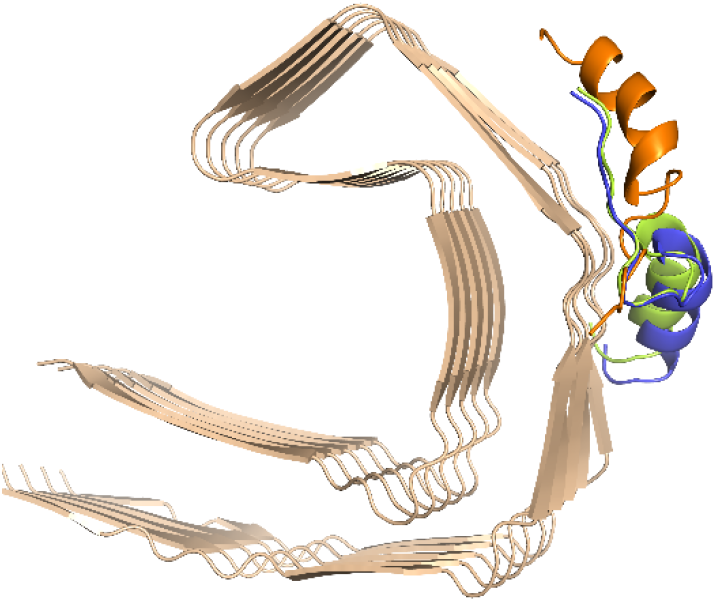
Docking result of Peptides p67 (limon0, p76 (orange), and p136 (blue) against Straight filaments (SFs) subunit.

## Conclusion

This study underscores the potential of peptide-based strategies for both treating Alzheimer’s disease by targeting tau aggregation. Through a combination of computational methods and peptide engineering, we have identified several peptides with high binding affinity for the PHF core, with p136 emerging as a standout candidate due to its substantial impact on PHF conformation and aggregation inhibition. The ability of p136 to induce significant structural disruptions and block additional PHF subunit binding highlights its potential as a therapeutic agent in AD. These insights pave the way for future research into peptide-based interventions for AD, emphasizing the need for continued exploration and validation in preclinical and clinical settings. Overall, this work contributes to the advancement of peptide-based diagnostic and therapeutic approaches, offering new hope for effective management of Alzheimer’s disease.

## Acknowledgements

None

